# Genetic Resilience of a Once Endangered Species, Tibetan Antelope (*Pantholops hodgsonii*)

**DOI:** 10.1101/628727

**Authors:** Yue Shi, Jiarui Chen, Jianping Su, Tongzuo Zhang, Samuel K. Wasser

## Abstract

Population reduction is generally assumed to reduce the population’s genetic diversity and hence its ability to adapt to environmental change. However, if life history traits that promote gene flow buffer populations from such impacts, conservation efforts should aim to maintain those traits in vulnerable species. Tibetan antelope (Pantholops hodgsonii) has experienced population reduction by 95% due to poaching during the 20^th^ century. We hypothesize that opportunities for gene flow provided by their sex-specific migration buffered their genetic diversity from the poaching impacts. We measured the mtDNA (control region, CR) and nuDNA (microsatellites or STRs) diversity, population differentiation, along with the change in effective population size (pre-poaching era vs. post-poaching era) and tested for a genetic bottleneck. Our results showed that Tibetan antelope maintained considerable genetic diversity in both mtDNA CR and STR markers (H_d_ = 0.9970 and H_obs_ = 0.8446, respectively), despite a marked reduction in post-poaching effective population size 368.9 (95% CI of 249.3 - 660.6) compared to the pre-poaching average (4.93×10^3^ - 4.17×10^4^). Post-poached populations also had low population structure and showed no evidence of a genetic bottleneck. Pairwise F_st_ values using CR haplotype frequencies were higher than those using STR allele frequencies, suggesting different degrees of gene flow mediated by females and males. This study suggests that the Tibetan antelope’s sex-specific migration buffered their loss of genetic diversity in the face of severe demographic decline. These findings highlight the importance of recognizing the traits likely to maintain genetic diversity and promoting conservation efforts that allow them to be exercised. For Tibetan antelope, this requires assuring that their migration routes remain unobstructed by growing human disturbances while continuing to enforce anti-poaching law enforcement efforts.

## Introduction

Biodiversity conservation is one of the most important priorities in conservation biology, where genetic diversity is an essential pillar. The need to conserve genetic diversity within populations is based on two arguments: (1) the necessity of genetic diversity for evolution in response to changing environments (Lacy, 1987; Morris, Austin, & Belov, 2012); (2) the expected correlation between genetic diversity and population fitness (Reed & Frankham, 2003) and links to the “extinction vortex” from inbreeding depression among fragmented populations (Frankham, 2005; Keller & Waller, 2002). Many natural populations have experienced severe demographic reduction due to rapid human population growth, overexploitation, environmental change, and habitat fragmentation. While it is generally assumed that population decline can drive the loss of genetic diversity, some species are able to maintain high genetic diversity even after a significant population crash (Busch, Waser, & DeWoody, 2007; Gonzalez-Suarez, 2010; Hailer et al., 2006; Kuo & Janzen, 2004; Lippé, Dumont, & Bernatchez, 2006; O’ Donnell, Richter, Dool, Monks, & Kerth, 2015). In such cases, life-history traits appear to buffer against the loss of genetic diversity in response to population declines (Hailer et al., 2006; Kuo & Janzen, 2004; Lippé et al., 2006). The life history of many migratory species offers great opportunities for gene flow via migration, introducing new alleles to the existing genetic diversity that might otherwise be lost from genetic drift (Busch et al., 2007; Frankham, 2015; Jangjoo, Matter, Roland, & Keyghobadi, 2016).

About one million Tibetan antelope (*Pantholops hodgsonii*) ranged across the Tibetan Plateau in the early 20^th^ century (Buzzard, Wong, & Zhang, 2012). However, Tibetan antelope populations were reduced to the brink of extinction at the end of the 20^th^ century by illegal poaching for their underfur, which was used to Shahtoosh shawls. Its population size reached a low of 50,000 individuals in 2003, declining by 95% relative to its size in 1950 (Leclerc, Bellard, Luque, & Courchamp, 2015). International conservation efforts successfully curbed the poaching through law enforcement and habitat protection, and by 2011, the number of Tibetan antelope was estimated to have increased to 200,000 individuals (Leclerc et al., 2015). The severe population reduction due to illegal poaching raised concerns regarding the genetic viability of Tibetan antelope populations and how it would affect their recovery.

Previous studies suggested that Tibetan antelope populations maintained high genetic variation with no signs of population structure (Ahmad et al., 2016; Du et al., 2016; H. Zhou, Li, Zhang, Yang, & Liu, 2007), although the genetic effects of the population crash on Tibetan antelope remain unclear. We hypothesize that the unique life history of Tibetan antelope may have buffered them against the loss of genetic diversity. Every summer, female Tibetan antelope from different wintering grounds migrate to the common calving ground to give birth, leave shortly after parturition and migrate back to their original wintering grounds with their newborn calves. However, not all females migrate back to their original wintering grounds (Buho et al., 2011). Male movements may also promote gene flow since there are no obvious geographic barriers on the Tibetan Plateau, although the movement of males remains unknown (Schaller, 1998). We assess the genetic diversity, population differentiation, population structure and effective population size of Tibetan antelope with maternal mtDNA (control region or CR) and bi-parental genetic markers (microsatellites or STRs). We predict that due to sex-specific migration, 1) there will be no obvious population structure; 2) differences in sex-specific migration could be reflected by population differentiation values using maternal markers vs. bi-parental marker; 3) Tibetan antelope populations are able to maintain high genetic diversity and show no signs of a population bottleneck.

## Materials and methods

### Ethics Statement

Tibetan antelope is listed in the Category I of the National Key Protected Wild Animal Species under the China’s Wild Animal Protection Law. In September 2016, Tibetan antelope was reclassified on the International Union for Conservation of Nature (IUCN) Red List from Endangered to Near Threatened due to their increased population size. Sample collection and field studies adhered to the Wild Animals Protection Law of the People’s Republic of China. Fresh scat samples were collected under IACUC protocol #2850-12. Dry skin samples and placenta samples were acquired with approval from the Forestry Department of Qinghai Province, China.

### Study Area and Sampling

We define all wintering grounds from which females that migrate to the same common calving ground as a deme. This unique migration pattern repeats itself for all the common calving grounds across the Tibetan Plateau (Schaller, 1998). The within-deme study focused on Zhuonai Lake in Kekexili Nature Reserve park (KKXL), Qinghai, China, which is the largest common calving ground for Tibetan antelope. Females from nearby wintering grounds migrate to Zhuonai Lake to give birth. Animals were observed with binoculars from a vigilance distance of ∼ 300m (Lian, Li, Zhou, & Yan, 2012) until they defecated and left the area. A total of 383 fresh scat samples were collected along with the date and GPS coordinates in ten different wintering grounds in KKXL (KKXL1 - KKXL10) and around the calving ground Zhuonai Lake (KKXL_ZNH). Samples were kept frozen until lab analyses. The among-deme study focused on three geographic populations of Tibetan antelope on the Tibetan Plateau, including the KKXL, examined above, along with Aerjin (AEJ) and Qiang Tang (QT) populations, using dry skin samples and placenta samples. The total sample size for the among-deme study was 141 (KKXL, N=69; AEJ, N=20; QT, N=52). See Figure 1 for sampling locations and Supplemental Table 1 for detailed sampling information.

**Figure 1.**
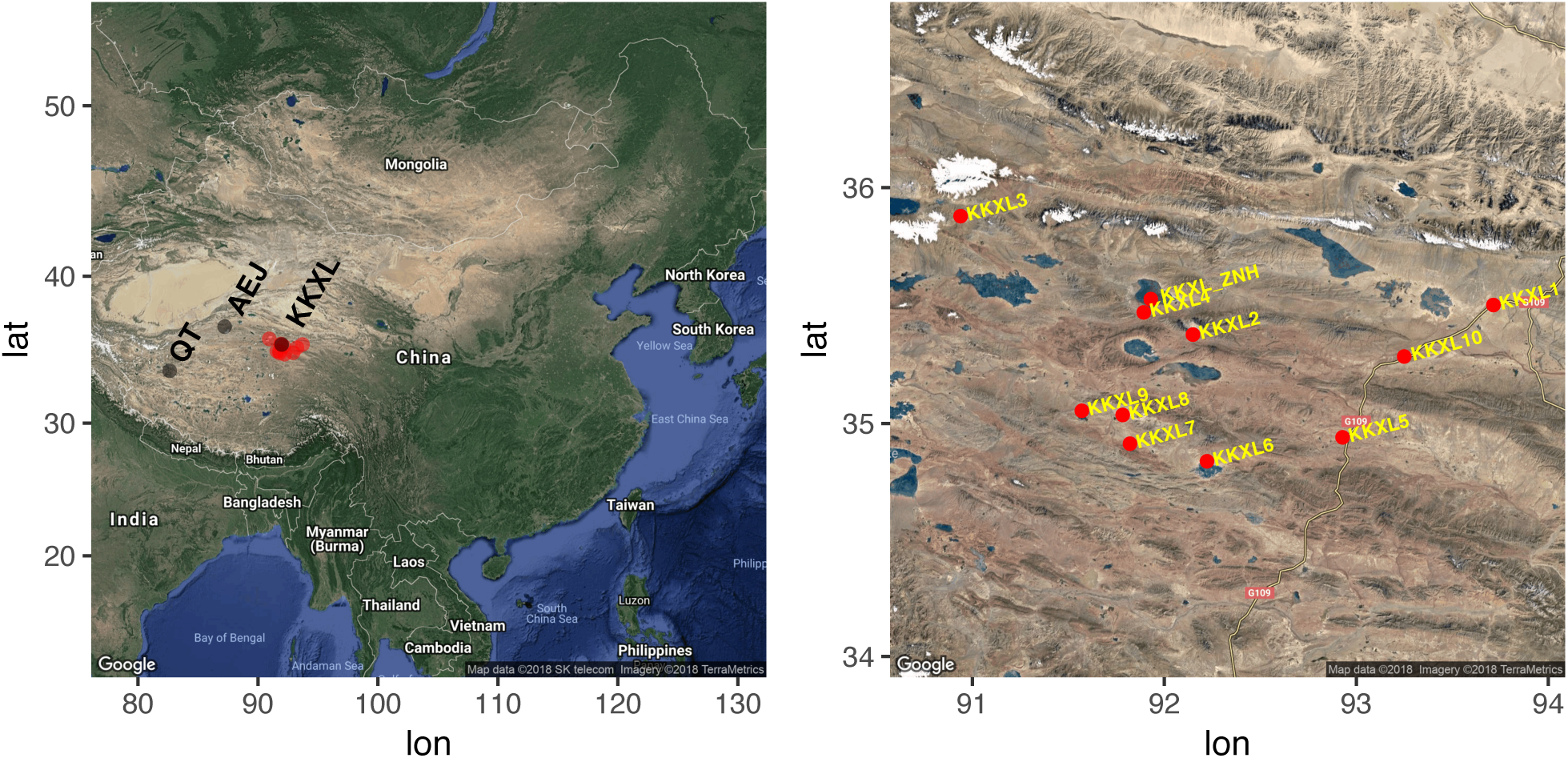
Sampling locations used for the among-demes study (left) and the within-demes study (right). Note: In the within-demes study, fresh scat samples were collected in ten different wintering grounds in KKXL (KKXL1 - KKXL10) and around the calving ground Zhuonai Lake (KKXL_ZNH). The among-deme study focused on three geographic populations of Chiru on the Tibetan Plateau, including the KKXL, examined above, along with Aerjin (AEJ) and Qiang Tang (QT) populations.

### DNA extraction and amplification

Fecal DNA was extracted and processed using the swabbing method (Wasser, Keim, Taper, & Lele, 2011). DNA in dry skins and placenta samples was extracted using the standard Phenol/Chloroform method (Strauss, 2001). mtDNA CR was amplified and sequenced in all samples using the forward primer DF (5’ ACCAGAGAAGGAGAACTCACTAACCT 3’) and the reverse primer DR (5’ AAGGCTGGGACCAAACCTAT 3’). PCR was conducted using the Qiagen Multiplex PCR kit (Qiagen Inc.,) with 0.5 μl 500 μg/mL of bovine serum albumin (BSA). We followed the recommended thermocycling conditions of the kit with an annealing temperature of 51° C.

Different sets of STR loci were used for within-deme and among-deme studies. The within-deme study used six STR loci denoted BM1824, MCM38, ILSTS005, MB066, BM1225, and BM4107 (see Supplemental Table 2A) with the 5’ end of forward primers fluorescently labeled with dyes 6-FAM or HEX. Annealing temperatures for PCR reactions for each locus were shown in Supplemental Table 2A. Fragment analysis was conducted on the ABI 3730 xl DNA Analyzer (Applied Biosystems). Alleles were scored with GeneMarker software (SoftGenetics, LLC.), and checked manually. We used duplicate fecal extracts and each extract underwent at least two independent PCR reactions to confirm allele profiles and guard against allelic dropout. Genotypes were classified as heterozygotes if both alleles were observed at least twice with no other alleles present, and homozygotes if only a single fragment was observed at least three times with no other alleles present. The among-deme study used seven STR loci denoted L01, L03, L04, ILSTS005, TGLA68, MCM38, and BM1341 (see Supplemental Table 2B). Amplification of these seven loci used the same protocol described in the within-deme study, but with a single extract amplified once per sample due to higher DNA yield.

### Mitochondrial CR sequence analyses

mtDNA CR sequences from both the within and among-deme studies were pooled together. All sequences were aligned using the software CLC Main Workbench (Qiagen, Inc). DnaSP v5.10.01 (Librado & Rozas, 2009) was used to determine the number of CR haplotypes (H), the number of segregating sites (S), haplotype diversity (*H*_*d*_) (Nei, 1978) and *H*_*d*_ standard deviation, nucleotide diversity (π) (Nei, 1978) and π standard deviation. We performed network analyses by constructing median-joining networks (Bandelt, Forster, & Rohl, 1999) on the control region haplotypes using the software PopART 1.7 (http://popart.otago.ac.nz).

### Nuclear STR analyses

Expected heterozygosity (*H*_exp_) observed heterozygosity (*H*_obs_), polymorphic information content (PIC) and estimated null allele frequency (*F*_null_) and combined probability of identity (PI, the probability of two independent samples having the save identical genotype by chance) were calculated using CERVUS v3.0.3 (Kalinowski, Taper, & Marshall, 2007). R packages *pegas* (Paradis, 2010) and *poppr* (Kamvar, Tabima, & Grünwald, 2014) were used to perform the exact test for Hardy-Weinberg equilibrium (HWE) and Linkage Equilibrium, respectively, using the Monte Carlo test with 1,000 iterations. Significance level was adjusted with sequential Bonferroni correction for multiple comparisons. Duplicate samples from the same genotypes were identified and excluded with the software CERVUS v3.0.3 (Kalinowski et al., 2007).

We evaluated population structure using Bayesian inference with the software STRUCTURE v2.2.3 considering an admixture model with correlated allele frequency (Pritchard, Stephens, & Donnelly, 2000). The individuals were assigned to possible genetic groups, K, varying from one to ten without a priori definition of populations. Twenty independent MCMC runs were carried out with 500,000 iterations following a burn-in period of 500,000 iterations for each value of the number of clusters (K). The best estimate of K was determined from both the likelihood of K and the ad hoc statistic delta K (Evanno et al. 2005). Genetic structure was also accessed through a discriminant analysis on principal components (DAPC) (Jombart, Devillard, & Balloux, 2010) implemented in the *adegenet* R package (Jombart, 2008).

### Population Differentiation Estimate

Pairwise *F*_*st*_ was assessed with 20,000 permutations in Arlequin v3.5.2 (Excoffier & Lischer, 2010) for mtDNA CR and STR data respectively. We performed an analysis of molecular variance (AMOVA) (Excoffier, Smouse, & Quattro, 1992) using Arlequin v3.5.2 to understand how genetic variation is partitioned. The significance of the proportion of variation at each category was obtained by MCMC test with 20,000 permutations. Isolation-by-distance (IBD) between Euclidean geographical distances and genetic distances (*F*_*st*_) were assessed using the Mantel test (Mantel 1967) with the *hierfstat* R package (Goudet, 2005) and 20,000 permutations. Significance values were adjusted for multiple comparisons using the Bonferroni correction (Rice 1989).

### Effective Population Size (*N*_*e*_) Estimation

We estimated historical (pre-poaching) and contemporary (post-poaching) effective population sizes of Tibetan antelope with the among-deme dataset. The historical effective population size *N*_*ef*_ of the mitochondrial genome was calculated from the CR region using the estimate of the female-specific theta (*θ*_*f*_ = 2*N*_*ef*_*μ*). *θ*_*f*_ estimate was derived from LAMARC (Kuhner, 2006) using Bayesian inference with 25 randomly selected samples, 10 initial search chains of 10,000 steps and 2 final chains of 1,000,000 iterations. A range of substitution rates (3.6 × 10^−10^ to 1.8 × 10^−8^ substitutions/site/gen) (Guo et al., 2006; Pesole, Gissi, De Chirico, & Saccone, 1999) in the CR region was used to reflect uncertainty in *μ*. The historical effective population size *N*_*e*_ was calculated based on the parameter theta *θ* = 4*N*_*e*_*μ* from LAMARC as with the estimate of *θ*_*f*_. *N*_*e*_ was estimated with 25 random samples using a range of STR mutation rates from 6 × 10^−5^ *to* 1.0 × 10^−3^ mutations/locus/generation (Crawford & Cuthbertson, 1996; Waples & Do, 2008) and the same running configuration as mtDNA CR sequences. The contemporary *N*_*e*_ was estimated using the linkage disequilibrium (LD) method in the software Ne Estimator v.2 (Do et al., 2014) with P_crit_ =0.01.

### Detection of Bottlenecks

We used four approaches to determine whether the overall Tibetan antelope population experienced a genetic bottleneck. First, we tested for allele frequency mode-shifts using BOTTLENECK v. 1.2.02 (Piry, Luikart, & Cornuet, 1999). Secondly, we tested for the presence of heterozygosity excess by using the one-tailed Wilcoxon signed rank test (Busch et al., 2007; Cornuet & Luikart, 1996) implemented in BOTTLENECK v. 1.2.02. Heterozygote excess was tested under all three STR mutation models: infinite alleles model (IAM), step-wise mutation model (SMM) and two-phase model (TPM). For TPM, we set p_s_=0.9 (the frequency of single-step mutations) and the variance of those mutations as 12 (Busch et al., 2007). Third, we calculated the M-ratio, the mean ratio of the number of alleles to the range in allele size, using the software M_P_VAL (Garza & Williamson, 2001). Critical values (M_c_) set at the lower 5% tail of the distribution were determined using the program CRITICAL_M. If the observed ratio is below M_c_, it can be assumed that the population has experienced a bottleneck (Garza & Williamson, 2001). To calculate M_c_, we estimated three TPM parameters: p_s_, Δ_g_ (the mean size of single-step changes) and pre-bottleneck *θ* = 4*N*_*e*_*μ*. We set p_s_ = 0.9, and Δ_g_ = 3.5. We varied *θ* from 0.01 to 500, encompassing a wide range of biologically plausible values. To ensure this range of *θ* values was relevant, we estimated *θ* using a common STR mutation rate *μ* (5.0 × 10^−4^ mutations/generation/locus) (Garza & Williamson, 2001) and *N*_*e*_ estimates from LAMARC. Lastly, we employed coalescent simulations with the Approximate Bayesian Computation (ABC) approach to infer past demographic history, as implemented in DIYABC v2.1.0 (Cornuet et al., 2014; Cornuet, Ravigne, & Estoup, 2010). Simulations were conducted with STR and mtDNA CR data separately. Only samples in the among-deme study were included. We compared two competing scenarios, scenario 1 with constant *N*_*e*_, and scenario 2 with population bottleneck (Supplemental Figure 2). The parameter settings and priors were shown in the Supplemental Table 3A & 3B and Supplemental method.

## Results

### Mitochondrial CR sequence analyses

The final alignment included 524 CR sequences of 1029 bp excluding insertions-deletions (indels). In total, we found 381 different haplotypes. All three geographical populations (KKXL, AEJ, QT) had high haplotype diversity (0.9890 - 1.000) and nucleotide diversity (0.0195 - 0.0241). The AEJ population had the largest standard deviation of haplotype diversity and nucleotide diversity, probably due to its small sample size (18 haplotypes in 20 sequences) (Table 1). The within-deme analysis of samples from 10 wintering locations in KKXL (KKXL1-KKXL10) revealed an overall pattern of haplotypes containing samples from multiple regions and no clusters with geographical affiliation (Supplemental Figure 1). In total, there were 17 haplotypes shared among 10 sampling locations from KKXL. The among-deme analysis of samples from KKXL, AEJ, and QT had a similar pattern (Supplemental Figure 1). There were two haplotypes shared between AEJ and QT, one haplotype shared between AEJ and KKXL, and one haplotype shared between QT and KKXL.

**Table 1.** Genetic diversity of Control Region according to geographical regions based on alignment of 524 sequences excluding indels (1029bp). Note: S = number of segregating sites (excluding sites with gaps/missing data); H_d_ = haplotype diversity; H_d_SD= standard deviation of H_d_; π = nucleotide diversity; πSD = standard deviation of π.

| Region | n | S | Haplotypes | $H_d$ | $H_dSD$ | $\pi$ | $\pi SD$ |
| --- | --- | --- | --- | --- | --- | --- | --- |
| KKXL<br>(within-deme) | 383 | 147 | 274 | 0.9960 | 0.0010 | 0.0206 | 0.0004 |
| KKXL | 69 | 119 | 68 | 1.0000 | 0.0030 | 0.0195 | 0.0011 |
| AEJ | 20 | 82 | 18 | 0.9890 | 0.0190 | 0.0241 | 0.0018 |
| QT | 52 | 95 | 48 | 0.9970 | 0.0040 | 0.0197 | 0.0011 |
| Total | 524 | 180 | 381 | 0.9970 | 0.0000 | 0.0203 | 0.0003 |

### Nuclear STR analyses

The set of STRs used in both within and among-deme studies revealed high power and accuracy. The combined probability of identity (PI) and sib identity (*P*_sib_) using all STRs in either study showed in Supplemental Table 4. In the within-deme study, the number of STR alleles per locus was 8-14, with an average of 11.167 (Table 2). All loci had high *H*_exp_ (0.704 - 0.875) and PIC (0.655 - 0.859), with mean *H*_exp_ of 0.7767 and PIC of 0.7451. Most loci were in HWE and Linkage Equilibrium across all populations after Bonferroni Correction. BM1824 violated HWE in KKXL10 population and MB066 violated HWE across all populations. BM1824 and MB066 showed signs of an excess of observed homozygote genotypes as suggested by relatively large positive *F*_null_ values (0.0410 and 0.0715, respectively) (Table 2). It is difficult to identify a null allele with certainty in the absence of a known parent-offspring relationship. Therefore, all loci were kept for further analyses.

**Table 2.** Genetic diversity and summary statistics of STR loci used in the study. A = number of alleles per locus; H_obs_ = observed heterozygosity; H_exp_ = expected heterozygosity; PIC = polymorphism information content; F_null_ = estimated frequency of null alleles (note: * p<0.05).

| Study | Loci | A | $H_{obs}$ | $H_{exp}$ | PIC | $F_{null}$ |
| --- | --- | --- | --- | --- | --- | --- |
| Within-deme study | BM1824 | 8 | 0.717 | 0.779 | 0.747 | 0.0410 |
|  | MCM38 | 9 | 0.739 | 0.704 | 0.655 | -0.0314 |
|  | ILSTS005 | 12 | 0.875 | 0.875 | 0.859 | -0.0012 |
|  | MB066 | 14 | 0.637 | 0.739 | 0.705 | 0.0715 |
|  | BM1225 | 12 | 0.768 | 0.785 | 0.762 | 0.0100 |
|  | BM4107 | 12 | 0.737 | 0.778 | 0.743 | 0.0270 |
| Among-deme study | L01 | 12 | 0.707 | 0.772 | 0.743 | 0.0508 |
|  | L03 | 24 | 0.887 | 0.928 | 0.921 | 0.0202 |
|  | L04 | 23 | 0.900 | 0.928 | 0.919 | 0.0141 |
|  | BM1341 | 15 | 0.858 | 0.885 | 0.872 | 0.0151 |
|  | ILSTS005 | 13 | 0.871 | 0.866 | 0.848 | -0.0040 |
|  | TGLA68 | 7 | 0.314 | 0.815 | 0.788 | 0.4450 |
|  | MCM38 | 9 | 0.702 | 0.716 | 0.679 | 0.0133 |

In the among-deme study, the number of STR alleles per locus was 7-24, with an average of 14.714 (Table 2). All loci had high *H*_obs_ except TGLA68 (0.314), and high PIC (0.679-0.921), with mean *H*_exp_ of 0.8446 and PIC of 0.8242. Most loci were in HWE across all populations after Bonferroni Correction. L03 violated HWE only in AEJ population and TGLA68 violated HWE across all populations. All loci were in Linkage Equilibrium except the pair of L04 and TGLA68, and L03 and L04. TGLA68 had a very high *F*_null_ value of 0.4450 (Table 2). TGLA68 was excluded from further analyses.

STRUCTURE analysis showed that the true K was equal to 1 for within and among-deme studies. DAPC analyses showed that there was no clear separation of 10 wintering locations within KKXL, and a high degree of overlap among the three geographical populations, KKXL, AEJ and QT (Figure 2).

**Figure 2.**
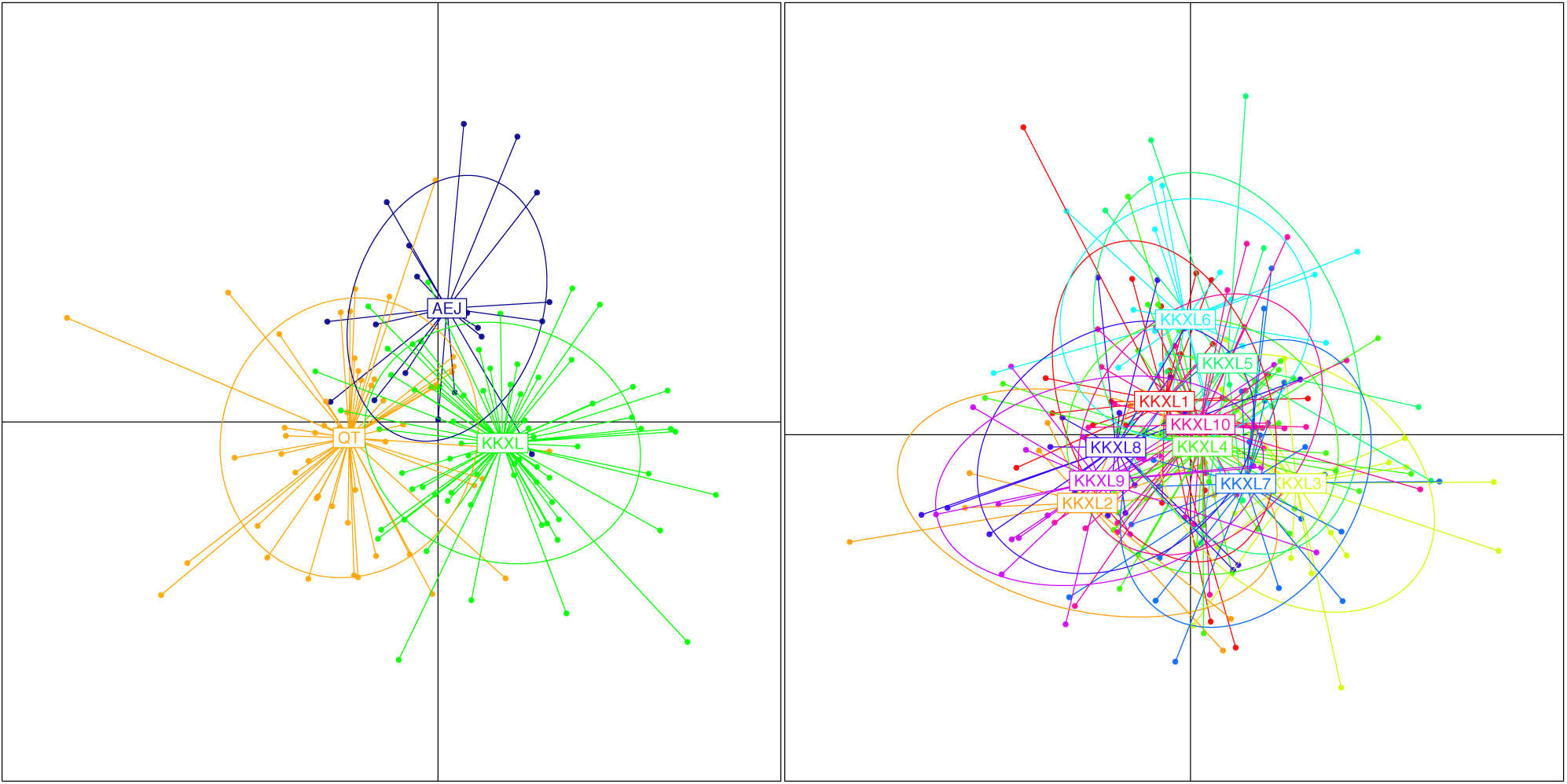
DAPC analyses with three geographical populations KKXL, AEJ and QT (left panel) and 10 sampling locations within KKXL (KKXL1-KKXL10, excluding KKXL_ZHN) (right panel) with discriminant function 1 on the x axis and discriminant 2 on the y axis.

### Population Differentiation Estimate

Population differentiation among geographic populations KKXL, AEJ and QT was low (Table 3), and only the *F*_*st*_ value calculated between AEJ and QT was significantly different from zero (Table 3). Pairwise *F*_*st*_ values were higher using the CR haplotype frequencies than the STR allele frequencies. Most genetic variation was attributed within populations instead of among populations (*p* values from AMOVA tests were 0.2308 and 0.3856 for CR haplotype frequency and STR allele frequency respectively). No significant correlation was detected between linearized genetic distances and geographic distance among KKXL, AEJ, and QT (*p* values from Mantel tests were 0.827 and 0.486 for CR haplotype frequency and STR allele frequency respectively).

**Table 3.** Pairwise F_st_ among populations KKXL, AEJ and QT calculated using control region haplotype frequencies (lower diagonal) and 6 microsatellite loci frequencies (six loci used in the large-scale study) (upper diagonal) with 20,000 permutations.

|  | AEJ | KKXL | QT |
| --- | --- | --- | --- |
| AEJ |  | 0.00424 | 0.00810* |
| KKXL | 0.01321 |  | 0.00536 |
| QT | 0.01560** | 0.00890 |  |

### Effective Population Size

The most probable estimate of *θ*_*f*_ from LAMARC was 0.084343 (95% CI 0.053719 - 0.134826) and the most probable estimate of *θ* from LAMARC was 9.989892 (95% CI 9.850306 - 10.01527). The long-term estimate of *N*_*ef*_ was in the range of 1.53 × 10^6^ - 1.79 × 10^8^, and the long-term estimate of *N*_*e*_ was in the range of 4.93 × 10^3^ - 4.17 × 10^4^. The contemporary *N*_*e*_ estimate was 368.9 (95% CI of 249.3 - 660.6).

### Bottleneck Analyses

The mode-shift test did not detect any evidence of a bottleneck (Table 4). Heterozygosity excess was detected only under the IAM mutation model (Table 4). While using Δ_g_=3.5, most M-ratio values were above the critical value thresholds except for the small pre-bottleneck *θ* values (0.01, 0.1 and 0.5) (See Table 4), which were very unlikely because historically Tibetan antelope had a large effective population size. All calculated M-ratios were above the suggested threshold value of 0.68 identified by Graza and Williamson (2001) for bottlenecked populations. ABC method also didn’t support the scenario of a population bottleneck. The best-supported model was constant population size model, with a posterior probability of 0.5258 (95% confidence interval CI: 0.5150 - 0.5366) for STR, and a posterior probability of 0.7142 (95% confidence interval CI: 0.7055 - 0.7229) for mtDNA CR sequences (Supplemental Figure 3). Analyses to estimate confidence in scenario choice indicated that type I (false-positive) and type II (false-negative) errors for the best-supported scenario (scenario 1 with constant *N*_*e*_) were high (0.438 and 0.389 for STR, 0.320 and 0.455 for mtDNA CR), suggesting low confidence in choosing the true scenario. Point estimate for *N*_*e*_ was 4.44 × 10^5^ (95% CI: 2.29 × 10^5^ - 9.45 × 10^5^) and 1.05 × 10^4^ (95% CI: 4.80 × 10^3^ - 1.93 × 10^4^) for mtDNA CR and STR respectively.

**Table 4.** Summary of the parameters and results for the M-ratio and BOTTLENECK analyses used to detect genetic bottleneck. NS=non-significant.

| $\theta$ | M-ratio | | Mode shift | BOTTLENECK | |
| --- | --- | --- | --- | --- | --- |
| | Mean M-ratio | $M_c$<br>($\Delta_g=3.5$) | | Mutation model | Heterozygote excess |
| 0.01 | 0.754 | 0.817 | NS |  |  |
| 0.1 | 0.754 | 0.817 |  |  |  |
| 0.5 | 0.754 | 0.780 |  |  |  |
| 1 | 0.754 | 0.749 |  | IAM | p=0.00781* |
| 5 | 0.754 | 0.712 |  | TPM | NS |
| 10 | 0.754 | 0.710 |  | SMM | NS |
| 50 | 0.754 | 0.715 |  |  |  |
| 100 | 0.754 | 0.705 |  |  |  |
| 500 | 0.754 | 0.640 |  |  |  |

## Discussion

This study tests the hypothesis that sex-specific migration can buffer populations from the impacts of population reduction with the case of migratory Tibetan antelope populations. Our results showed that 1) Tibetan antelope maintained high genetic diversity in both mtDNA CR and STR markers after a historical population decline; 2) No population bottleneck was detected; 3) There was no obvious population structure among three geographical populations, which is a sign of high gene flow among populations. Males are likely to play a bigger role in gene flow than females since pairwise *F*_*st*_ values were higher using the maternal CR haplotype frequencies than the biparental STR allele frequencies. This study suggests that the Tibetan antelope’s sex-specific migration reduces their loss of genetic diversity in the face of severe demographic decline. However, Tibetan antelope has not fully recovered from poaching in terms of effective population size. There is a marked reduction in post-poaching effective population size 368.9 (95% CI of 249.3 - 660.6) compared to the pre-poaching average (4.93 × 10^3^ - 4.17 × 10^4^).

### Tibetan antelope maintained high genetic diversity after a historical population decline

Overall, CR haplotype diversity (*H*_d_) was 0.9975 and π was 0.02026. The mean *H*_exp_ of STR loci was 0.7767 and 0.8446 for the within and among-deme studies respectively. Our results are consistent with previous studies (Du et al., 2016; F. Zhang, Jiang, Xu, Zeng, & Li, 2013). At the species level, the genetic diversity of Tibetan antelope was higher than other endangered ungulate species, such as Roan antelope (*Hippotragus equinus*: size 401bp; *H*_d_ 0.6969; π 0.0190; *H*_exp_ 0.46), Saiga antelope (*Saiga tatarica*: size 277 bp; *H*_d_ 0.785; π 0.0139) and Kashmir red deer (*Cervus elaphus hanglu*: size 454 bp; *H*_d_ 0.589; π 0.008; *H*_exp_ 0.66) (Alpers, Van Vuuren, Arctander, & Robinson, 2004; Campos et al., 2010; Mukesh, Kumar, Sharma, Shukla, & Sathyakumar, 2015; Singh, Grachev, Bekenov, & Milner-Gulland, 2010).

### No population bottleneck was detected

Neither the mode-shift, heterozygosity excess, M-ratio nor ABC method revealed strong evidence of a population bottleneck. The mode-shift test did not detect any evidence of a bottleneck. Heterozygosity excess was detected only under the IAM model. IAM is prone to incorrectly detect heterozygosity excess in non-bottlenecked populations. Therefore, to be statistically conservative, one should use the SMM or TPM when analyzing STR data to test for recent bottlenecks (Luikart and Cornuet, 1998). The M-ratio approach detected bottleneck signatures, but only under extreme conditions (very small 7 values), suggesting a weak signal, if any. The model with constant population size has higher support over the model with population bottleneck based on ABC method, though it has high Type I and II errors.

### No obvious population structure was detected among three geographical populations

Despite large-scale sampling efforts, phylogenetic analysis with Bayesian inference, haplotype network analysis of the CR region, STRUCURE and DAPC analyses of STR loci revealed no obvious geographic structure for Tibetan antelope in AEJ, QT, and KKXL populations. The Mantel test detected no IBD pattern with neither mtDNA CR nor STR loci. This finding suggests historically wide gene flow, which is consistent with results of previous studies (F. Zhang et al., 2013; H. Zhou et al., 2007).

There are no obvious geographic barriers blocking population exchange on the plateau. During the course of female Tibetan antelope migration, it is possible that a number of females from one population translocate to another. This would promote gene exchange between populations of different localities, which is reflected by the shared haplotypes among different populations (Supplemental Figure 1). However, males are likely to play a bigger role in gene flow since pairwise *F*_*st*_ values were higher using the maternal CR haplotype frequencies than the biparental STR allele frequencies. Tibetan antelope have a harem polygyny mating system, in which a male generally mates with most or all of the females in his harem (5-10 females) during the breeding season. Mate competition is an important driver explaining the spatial movement of males among populations during the breeding season. Breeding dispersal is not restricted to young males. It also occurs among prime-aged individuals and even among harem holders (Jarnemo, 2011; Richard, White, & Côté, 2014).

### Tibetan antelope has not fully recovered from poaching yet in terms of effective population size

In 2003, the estimation of Tibetan antelope population size reached the lowest number of 50,000 individuals. Since then, the Tibetan antelope population has begun to recover, with about 200,000 individuals currently (Leclerc et al., 2015). Their protection status has been changed from “endangered” to “near-threatened” by IUCN. However, our effective population size comparison analyses suggest that Tibetan antelope has not yet fully recovered. Their contemporary *N*_*e*_ estimate is 368.9 (95% CI of 249.3 - 660.6), which is markedly lower than their long-term *N*_*e*_ average (4.93 × 10^3^ - 4.17 × 10^4^). Long-term *N*_*e*_ estimate with ABC method using STR loci is 1.05 × 10^4^ (95% CI: 4.80 × 10^3^ - 1.93 × 10^4^), which mostly agrees with the estimate with LAMARC method.

The contemporary *N*_*e*_ estimate has a wide confidence interval (95% CI of 249.3 - 660.6). In the LD method implemented in Ne Estimator v.2, CI of *N*_*e*_ is an increasing function of *N*_*e*_ (Posada & Crandall, 2001; Waples & Do, 2010). Like all the other genetic methods for estimating contemporary *N*_*e*_, LD method is most powerful with small populations and has difficulty distinguishing large populations from infinite ones. However, it should provide a useful lower bound for *N*_*e*_, which can be important in conservation biology where a major concern is avoidance and early detection of population bottlenecks (Bandelt et al., 1999; Waples & Do, 2010).

Surprisingly, the mtDNA CR-based estimate of *N*_*ef*_ (1.53 × 10^6^ - 1.79 × 10^8^) was larger than the STR-based estimate *N*_*e*_ (4.93 × 10^3^ - 4.17 × 10^4^). Long-term *N*_*e*_ estimate with ABC method using mtDNA CR sequences is 4.44 × 10^5^ (95% CI: 2.29 × 10^5^ - 9.45 × 10^5^), which is lower than the estimate with LAMARC, but still larger than the STR-based estimate. In theory, the mitochondrial genome has an effective population size one quarter that of an average nuclear locus because of the different inheritance modes of nuclear and mtDNA, as well as the haploid nature of the mitochondrial genome. The observed disparity between mtDNA and STR-based estimates could result from the Tibetan antelope’s mating system, historical demography, mutation or all three combined. Tibetan antelope have a harem polygyny mating system, as mentioned above. Non-independent mating paring has a large effect when there is intense male-male competition for reproduction in a harem social system and reduces *N*_*e*_ for wholly or paternally inherited components of genome (Evans & Charlesworth, 2013; Nunney, 1996). Historical events may also explain the apparent discord between mtDNA and STR-based of longterm population size. For instance, if the historical source populations that contributed to the origin of the contemporary population were isolated from each other but with male-biased dispersal, then the populations would more rapidly diverge at mtDNA loci than nuclear loci because mtDNA is matrilineally inherited. A secondary contact of these isolates would merge a relatively homogenous pool of nuclear genes, but mtDNA lineages would remain differentiated. This evolutionary scenario is supported by our pairwise *F*_*st*_ results (Table 3) that Pairwise *F*_*st*_ values were higher using the CR haplotype frequencies than the STR allele frequencies. *F*_*ef*_ may reflect the collective genetic diversity of all source populations, which would possibly contribute to the reversal in expected sizes for *N*_*e*_ and *N*_*ef*_. Another possibility is that size homoplasy of STRs might have obscured the signal on the historical origin of the study population. Some STR alleles can be identical in size but may not identical by descent due to convergent mutations. Size homoplasy is especially problematic in large populations (Estoup, Jarne, & Cornuet, 2002), such as Tibetan antelope population. Thus, STR-derived *N*_*e*_ estimates may not reflect the composite origin of these populations as well as *N*_*ef*_. Regardless of the ultimate cause of the discord between *N*_*e*_ and *N*_*ef*_, all estimators indicate that the historical population size of Tibetan antelope is very large.

### How did Tibetan antelope maintain such high genetic diversity despite a massive population decline?

We expected the Tibetan antelope to have suffered a serious population bottleneck from the near 95% decline in the original population due to poaching. However, we found no evidence of such an event by any of the methods used in the study. Tibetan antelope still maintains high genetic diversity.

The contemporary *N*_*e*_ of Tibetan antelope estimate was 368.9, with a wide 95% confidence interval of 249.3 - 660.6, which is large enough to preclude losses of neutral genetic diversity. Small populations caused by massive population reduction are at risk of extinction vortex due to demographic stochasticity and random genetic drift. The genetic pool is seriously impaired when the effective population size reduces below the threshold of 50, according to the 50/500 rule, which proposes that a population would need to begin with at least a population size of 50 individuals to ensure enough initial genetic variation, and maintain a population size of 500 to keep the levels of genetic variation over time (Franklin, 1980).

It is also possible that insufficient time has passed since the start of population reduction to markedly reduce Tibetan antelope genetic diversity. It is hard to detect a recent population bottleneck, as reflected by the high type I and II error in DIYABC analyses. According to the coalescent theory (Crow & Kimura, 1970), *H*_*t*_/*H*_*o*_ (the ratio between heterozygosity at generation t vs. generation 0) depends on (1 − 1/2N)^t^. For populations with fluctuating population size from generation to generation, one should replace N with harmonic mean of the generation-specific effective sizes *N_e_*\* If we assume that the year 1950 is time 0 and *N_e_*\* = 249 (we chose the lower bound of contemporary effective population size as *N_e_*\* to be conservative), by 2016 *H*_*t*_/*H*_*o*_ would be 0.936 (assuming generation time is 2 years). Thus, we would not expect to see a significant loss of genetic diversity by genetic drift.

The high genetic variability in Tibetan antelope population after a population crash likely reflects the effect of admixture. A few immigrants entering a population each generation can counteract the effects of genetic drift and obscure any genetic signature of this population’s decline. Increasing human activities on the Tibetan Plateau is now threatening this once-paradise for wildlife. Habitat fragmentation, such as fencing and road construction, might affect gene flow among populations, leading to local extinction that could impede Tibetan antelope population’s recovery (X. Su et al., 2015; Zhuge, Li, Zhang, Gao, & Xu, 2014).

### Future Research Recommendations

We acknowledge that the few loci used in this study provide limited resolution and considerable uncertainty for demographic inference. Each locus on the genome has a particular genealogy describing its history. Genetic data using a few loci only provides a few numbers of possible genealogies for the underlying demography. The only way to reduce the uncertainty of the demographic model is to sample throughout the entire genome, which will contain a wealth of information for demographic inference.

When the changes in genetic diversity are assessed for endangered populations, it is essential to establish a baseline representing the pre-poaching conditions (Matocq & Villablanca, 2001). The use of archival (e.g. museum samples) specimens may allow for a powerful test of loss in genetic diversity over time. If archival reference samples represent genetic variation found in a population prior to the events leading to endangered status (e.g. samples from the pre-poaching era), such samples would provide the most appropriate reference against which to measure current levels of diversity (Bouzat, Lewin, & Paige, 1998).

## Acknowledgments

We thank Hyeon Jeong Kim, Adam Leaché, Chad Klumb for valuable comments that greatly improved this manuscript. We thank Rebecca Booth for lab work advice. We are thankful to Lei Wang, Jundong Yang and Jie Yang for the help with sample collection and lab work. We thank Kekexili Natural Nature Reserve Administration for assistance in the fieldwork. The study was funded by 1) Biodiversity Study in Kekexili, Department of Housing and Urban-Rural Development of Qinghai Province, China; 2) Construction Fund for Qinghai Key Laboratories (2017-ZJ-Y23); 3) The Strategic Priority Research Program of the Chinese Academy of Sciences (XDA23060602, XDA2002030302); 4) Wingfield-Ramenofsky Award, University of Washington. Yue Shi was funded by China Scholarship Council (CSC) Graduate Research Fellowship, Fritz/Boeing International Research Fellowship and WRF Hall Fellowship from the University of Washington.

## Data Accessibility

After peer review and prior to final publication, mtDNA sequences will be deposited in GenBank and microsatellite genotype dataset will be archived at Dryad.

## Conflict of interests

The authors declare no conflict of interests.

## Author Contributions

Y.S., S.K.W., and J.P.S. designed the project. J.R.C., Y.S., T.Z.Z. and J.P.S. conducted sample collection in the field. Y.S. and J.R.C. performed the lab experiment. J.P.S. oversaw the study and offered insight into data analysis and interpretation. Y.S. wrote the manuscript, and all other authors provided comments and approved the final version.

## Notes

#### Summary of Updates

Contact info updated; Acknowledgments updated; Supplemental files updated.

