## Supplemental for "Genetic Resilience of a Once Endangered Species, Tibetan Antelope (*Pantholops hodgsonii*)"

**Supplemental Table 1** Sampling location Information

| Study | Sampling Site ID | Latitude | Longitude | Sample Type | Sampling Time | Sample Size |
|---|---|---|---|---|---|---|
| Within-deme | KKXL1 | 35.5035 | 93.7141 | Feces | May, 2015/ April, 2016 | 24 |
| | KKXL 2 | 35.3782 | 92.1489 | Feces | May, 2015 | 10 |
| | KKXL 3 | 35.8795 | 90.9389 | Feces | May, 2015 | 23 |
| | KKXL 4 | 35.4723 | 91.8928 | Feces | May, 2015 | 44 |
| | KKXL 5 | 34.9406 | 92.9283 | Feces | Apr, 2016 | 15 |
| | KKXL 6 | 34.8383 | 92.2222 | Feces | Apr, 2016 | 20 |
| | KKXL 7 | 34.9137 | 91.8211 | Feces | Apr, 2016 | 22 |
| | KKXL 8 | 35.0362 | 91.7831 | Feces | Apr, 2016 | 20 |
| | KKXL 9 | 35.0539 | 91.5703 | Feces | Apr, 2016 | 20 |
| | KKXL 10 | 35.2850 | 93.2494 | Feces | May, 2015/ Apr, 2016 | 36 |
| | KKXL_ZNH | 35.5285 | 91.9295 | Feces | Jul, 2015 | 149 |
| Among-deme | QT | 33.6961 | 82.6392 | Dry skin | Sep, 2013 | 52 |
| | AEJ | 36.7015 | 87.2235 | Placenta | Jul, 2014 | 20 |
| | KKXL | 35.4983 | 91.9769 | Placenta | Jul, 2014 | 69 |

**Supplemental Table 2A** Six polymorphic microsatellite loci selected for fecal samples in the within-deme study

| Loci | Primer (5'-3') | Ta (°C) | Multiplex Set | Reference |
|---|---|---|---|---|
| BM1824 | F: GAGCAAGGTGTTTTTCCAATC<br>R: CATTCTCCAACTGCTTCCTTG | 58 | 1 | (Bishop et al., 1994) |
| MCM38 | F: TGGTGAATGGTGCTCTCATACCAG<br>R: CAGCCAGCAGCCTCTAAAGGAC | 58* | 2 | (H. Zhou, Li, Zhang, Yang, & Liu, 2007) |
| ILSTS005 | F: GGAAGCAATGAAATCTATAGCC<br>R: TGTTCTGTGAGTTTGTAAGC | 55 | 3 | (Brezinsky, Kemp, & Teale, 1993) |
| MB066 | F: ATCTGCCTGAAGCCAGTCAC<br>R: GGTTTCCTGCACCTGCATGA | 56 | 4 | (H. Zhou et al., 2007) |
| BM1225 | F: TTTCTCAACAGAGGTGTCCAC<br>R: ACCCCTATCACCATGCTCTG | 56 | 4 | (Bishop et al., 1994) |
| BM4107 | F: AGCCCCTGCTATTGTGTGAG<br>R: ATAGGCTTTGCATTGTTCAGG | 56 | 5 | (Bishop et al., 1994) |

Note: All loci except for MCM38 were amplified as following: 95 °C (15 min), then 40 cycles at 94°C (30s) / Ta°C (90s) / 72°C (60s), and a final extension at 60°C for 30 min. MB066 and BM1225 were amplified in a duplex PCR. *: MCM38 was amplified in a touchdown PCR. 95 °C (15 min), then 12 cycles at 94°C (30s) / 70°C (90s) (with decrement of 1°C per cycle) / 72°C (60s), 28 cycles at 94°C (30s) / 58°C (90s)/ 72°C (60s) and a final extension at 60°C for 30 min.

**Supplemental Table 2B** Seven polymorphic microsatellite loci selected for dry skin and placenta samples in the among-deme study

| Loci | Primer (5'-3') | Ta (°C) | Reference |
|---|---|---|---|
| L01 | F:TCTTGTGATCTCTTCCAGTAGAG<br>R:CGTCAGGCAATGAAGGTAG | 54 | (Zhou et al., 2014) |
| L03 | F:CTGACTTCTTTCTCCCTACGA<br>R:CAACCACTTTTGGATTCACAG | 54 | (Zhou et al., 2014) |
| L04 | F:CAAGGGATCATTTCAATGCT<br>R:TGTTCTGTGAGTTTGTAAGC | 58.5 | (Zhou et al., 2014) |
| ILSTS005 | F:GGAAGCAATGAAATCTATAGCC<br>R:TGTTCTGTGAGTTTGTAAGC | 58 | (Brezinsky et al., 1993) |
| TGLA68 | F:ATCTTACTTACCTTCTCAGCGCT<br>R:GGGACAAAATTTTACATATACACTT | 59 | (H. Zhou et al., 2007) |
| MCM38 | F:TGGTGAATGGTGCTCTCATACCAG<br>R:CAGCCAGCAGCCTCTAAAGGAC | 58 | (H. Zhou et al., 2007) |
| BM1341 | F:CCTACCTACTGCACAGTTTTGC<br>R:CTCCCATATAAGTTACCCACCC | 60 | (H. Zhou et al., 2007) |

Note: All loci were amplified as following: as following: 95 °C (15 min), then 40 cycles at 94°C (30s) / Ta°C (90s) / 72°C (60s), and a final extension at 60°C for 30 min in singlex PCR.

**Supplemental Table 3A** Demographic parameters used for the Approximate Bayesian Computation (ABC) models of constant population size, population bottleneck with among-deme microsatellite dataset. N – Effective population size; Na – Ancestral population size; Nc – Contemporary population size; t: time of population bottleneck.

| Parameter | Prior (uniform distribution) |
|---|---|
| Na | 10-20,000 |
| Nb | 10-20,000 |
| Nc (Nc<=Na) | 10-20,000 |
| Tb (in generations) | 0-100 |

**Supplemental Table 3B** Demographic parameters used for the Approximate Bayesian Computation (ABC) models of constant population size, population bottleneck with mtDNA CR sequences. N – Effective population size; Na – Ancestral population size; Nc – Contemporary population size; t: time of population bottleneck.

| Parameter | Prior (uniform distribution) |
|---|---|
| Na | 10-1,000,000 |
| Nb | 10-1,000,000 |
| Nc (Nc<=Na) | 10-1,000,000 |
| Tb (in generations) | 0-100,000 |

**Supplemental Table 4** Combined non-exclusion probability for STR loci used in studies. PI: combined non-exclusion probability for identity; $P_{sib}$: combined non-exclusion probability for sib identity.

| Study | PI | $P_{sib}$ |
|---|---|---|
| within-deme | $5.28 \times 10^{-11}$ | $2.90 \times -7$ |
| among-deme | $5.10 \times 10^{-4}$ | $3.40 \times 10^{-3}$ |

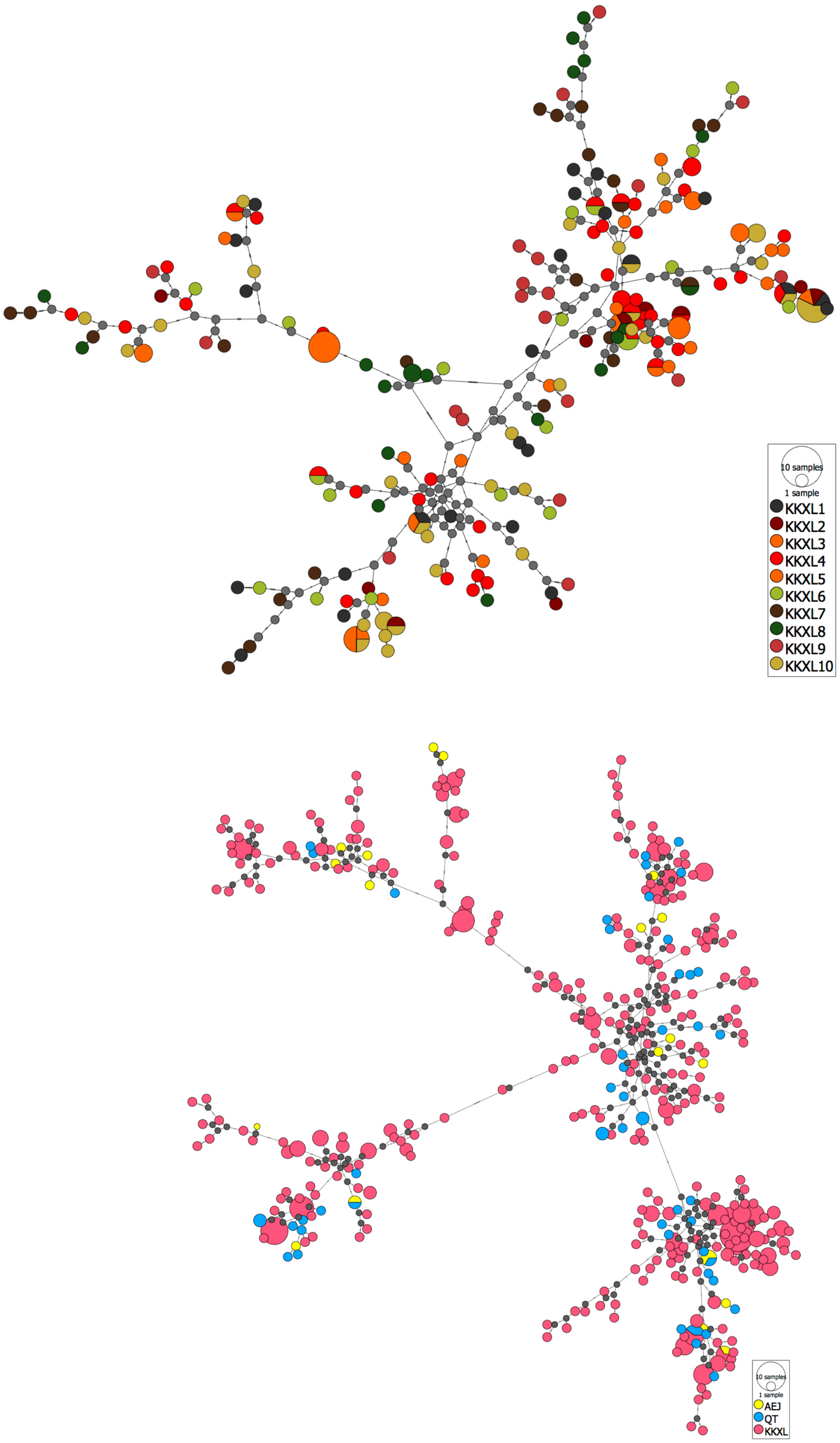


**Supplemental Figure 1** Median-joining network analysis based on 1029 bp control region haplotypes (excluding indels). Top panel shows 190 haplotypes from 10 wintering locations within KKXL (excluding KKXL_ZHN). Bottom panel shows 381 haplotypes from three geographical populations AEJ, QT, and KKXL.

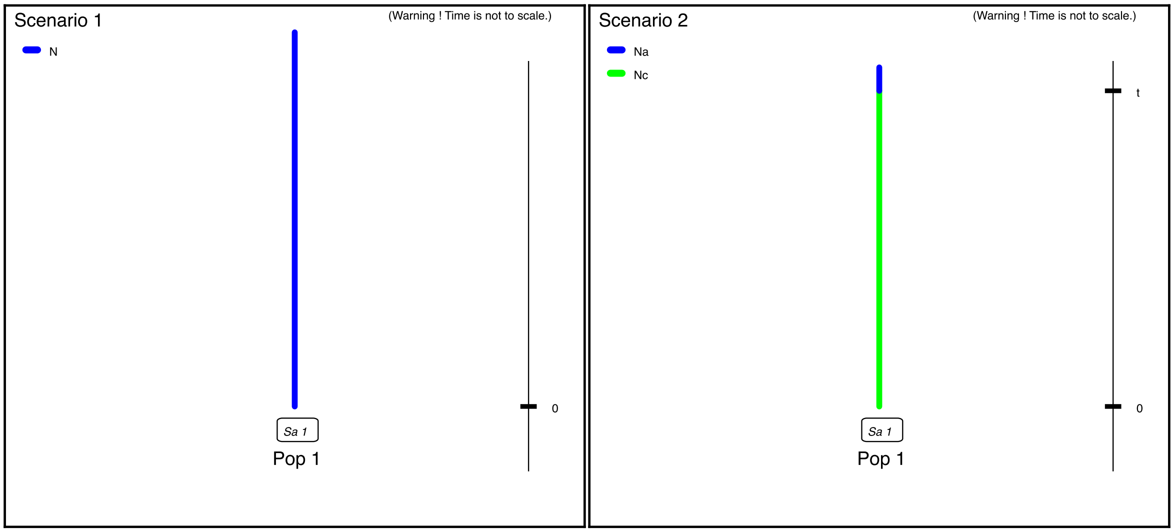


**Supplemental Figure 2** Demographic scenarios for comparison used for ABC simulations. Scenario 1 is for constant population size and Scenario 2 is for population bottleneck.

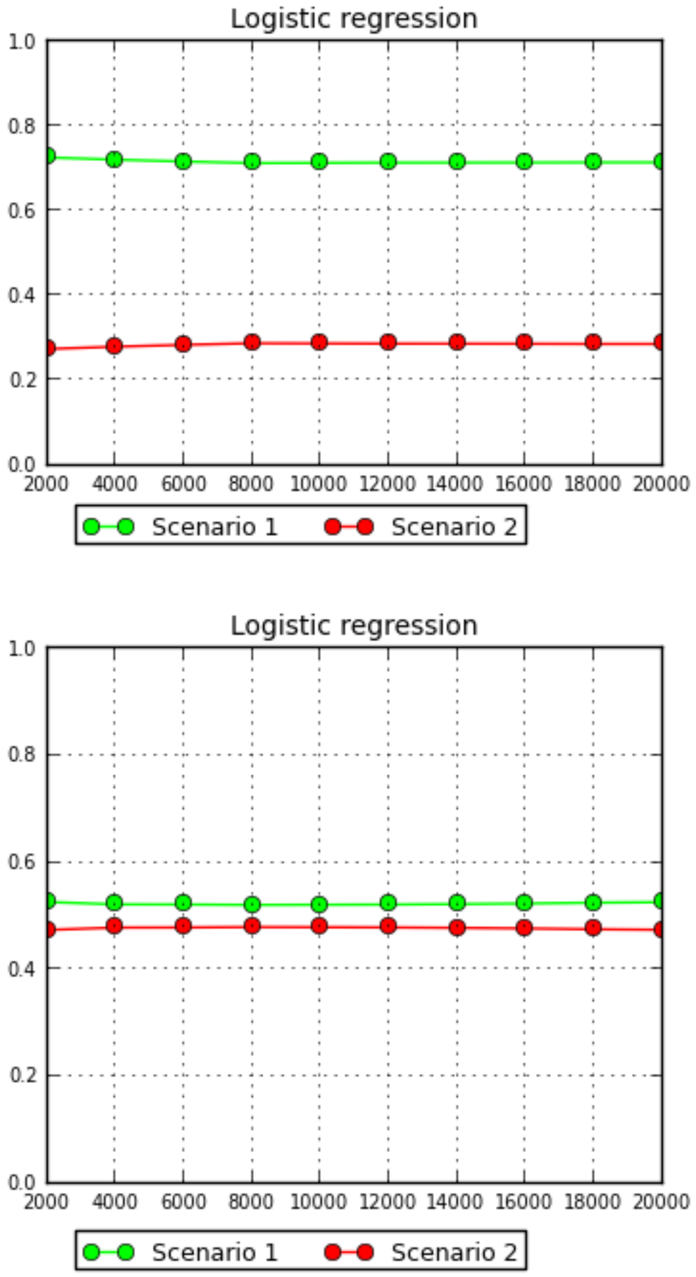


**Supplemental Figure 3** Posterior probability of models in comparison with logistic regression implemented in DIYABC. Note: Scenario 1 is for constant population size and Scenario 2 is for population bottleneck. Top panel was based on simulation with mtDNA CR sequences and bottle panel was based on simulation with STR loci.

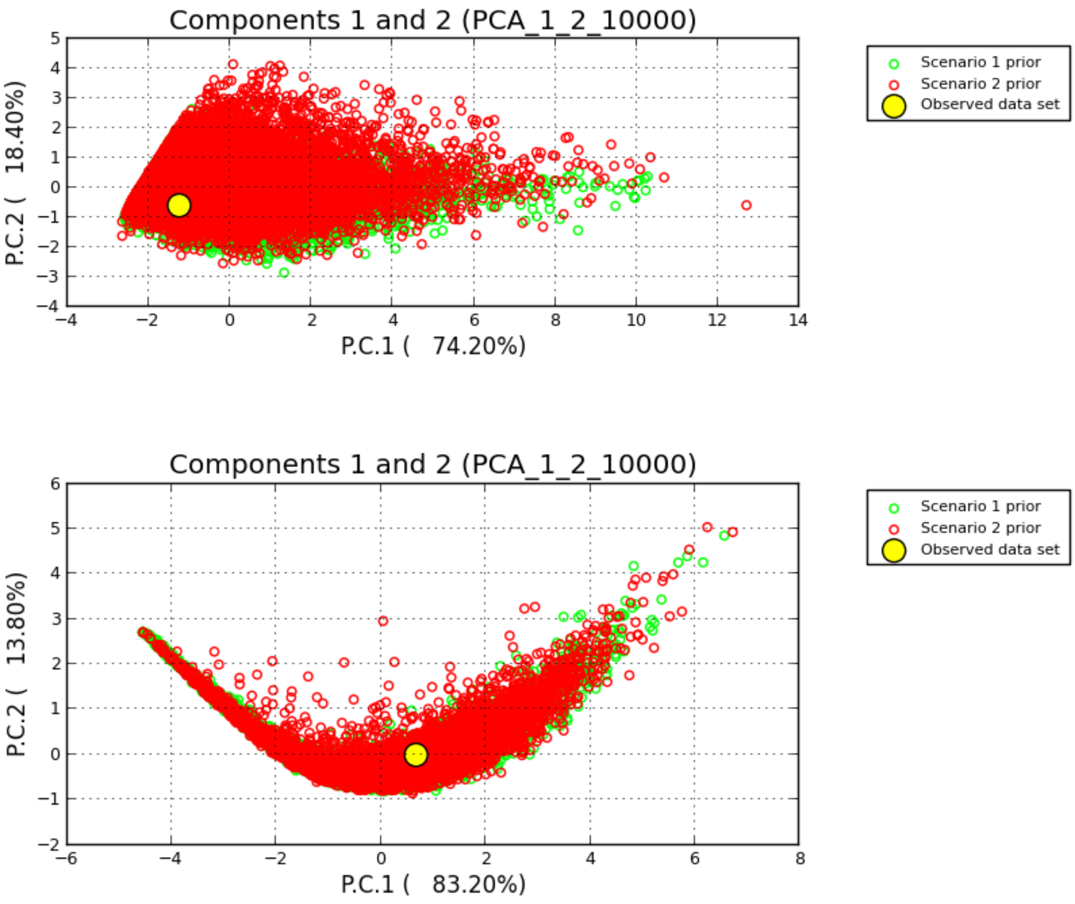


**Supplemental Figure 4** Pre-evaluation of scenario-prior combination suitability with PCA analyses implemented in DIYABC. Observed dataset was placed within the 1% of simulated data sets. This suggests that the selected summary statistics and priors were suited for the models. Note: Scenario 1 is for constant population size and Scenario 2 is for population bottleneck. Top panel was based on simulation with mtDNA CR sequences and bottle panel was based on simulation with STR loci.

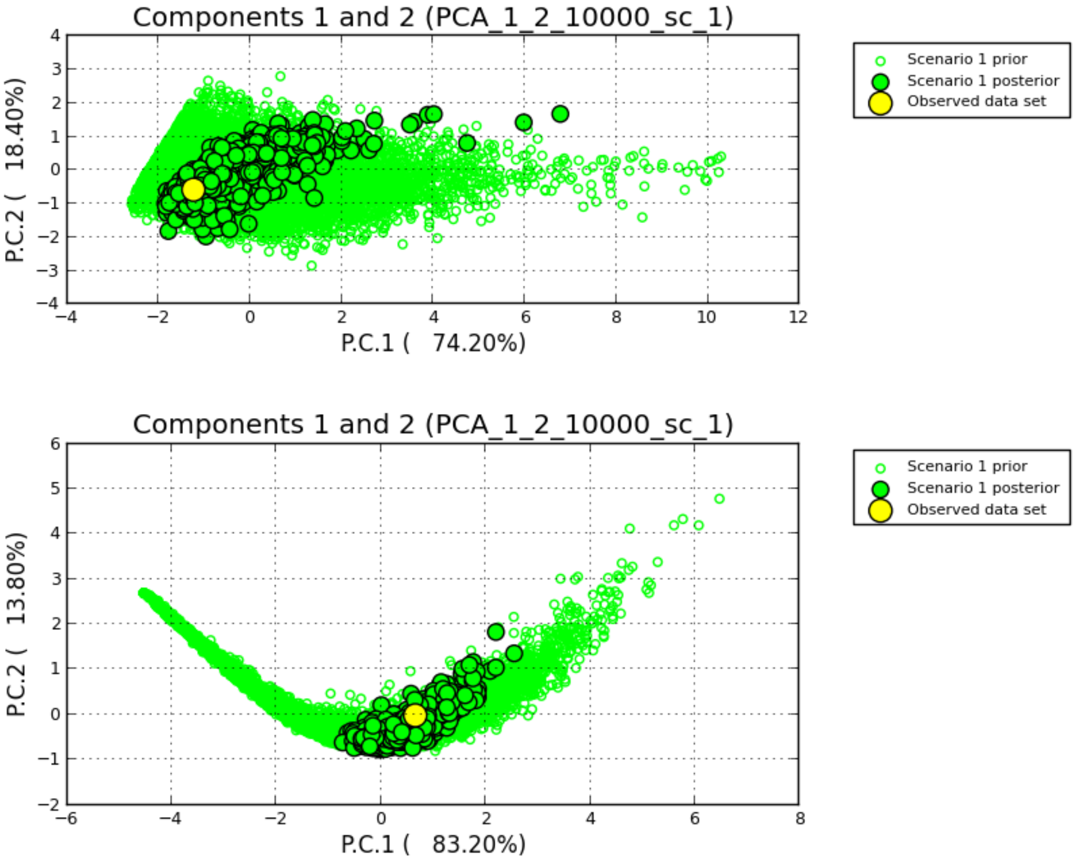


**Supplemental Figure 5** Model checking to assess the goodness-of-fit for the best-supported scenario (Scenario 1 with constant population size) implemented in DIYABC. Model checking showed a large cluster of simulated data from the prior and a small cluster of data from the posterior predictive distribution with the observed data set placed within both, suggesting the model/posterior for scenario 1 provided a good fit to the observed data. Note: Top panel was based on simulation with mtDNA CR sequences and bottle panel was based on simulation with STR loci.

### Supplemental method --- DIYABC methods

Larger demographic and temporal prior ranges were given for mtDNA CR sequences due to their larger $N_e$ estimates from initials tests. Because no genetic structure was found in Chiru populations, both scenarios were simulated under the framework of a single population. We used a Generalized Stepwise Mutation (GSM) model with a mean STR mutation rate of $10^{-5}$ to $10^{-3}$ drawn from a uniform distribution. We set the shape to 0 so all individual loci took the same values (=mean). The prior for the mutation rate in mtDNA CR sequence was set to draw from a gamma distribution with mean $2.109 \times 10^{-7}$with mutation model as Hasegawa-Kishino-Yano or HKY (1985). All other default settings remained in place. We chose the following as summary statistics: 1) for mtDNA CR sequences: number of segregating sites, mean of pairwise differences, variance of pairwise differences and Tajima's D; 2) for STRs: mean number of alleles. Mean genic diversity and mean size variance. A total of $2 \times 10^6$ data sets were simulated and summary statistics per scenario were calculated on each simulation, with roughly equal representation of each scenario in the reference table. We used Principal Component Analysis (PCA) in the "pre-evaluate" option to initially evaluate scenario-prior combinations. To compare scenarios, we computed the posterior probability of each of them by performing a logistic regression based on 1% of the simulated data closest to observed data. The scenario with the higher posterior probability was selected as the best. For the best-supported scenario, we performed the option "model check" to assess the goodness-of-fit. A good fitting model should produce a cloud of data points simulated from the priors, on top of which lies the observed data within a smaller cloud of datasets from the posterior predictions. To assess confidence in scenario choice, we calculated type I and II error rates from 20,000 pseudo-observed datasets (PODs) for the "logistic approach. We measured the proportion of times that the best-fit scenario had the highest posterior probability compared the competing scenario out of 1000 requested PODs. Point estimates for demographic and temporal parameters were obtained by local linear regression on the 1% of simulated data sets closest to the observed dataset for the best-supported scenarios.
